# Automated workflow for BioID improves reproducibility and identification of protein-protein interactions

**DOI:** 10.1101/2023.09.08.556804

**Authors:** Emilio Cirri, Hannah Knaudt, Domenico Di Fraia, Nadine Pömpner, Norman Rahnis, Ivonne Heinze, Alessandro Ori, Therese Dau

## Abstract

Proximity dependent biotinylation is an important method to study protein-protein interactions in cells, for which an expanding number of applications has been proposed. The laborious and time consuming sample processing has limited project sizes so far. Here, we introduce an automated workflow on a liquid handler to process up to 96 samples at a time. The automation does not only allow higher sample numbers to be processed in parallel, but also improves reproducibility and lowers the minimal sample input. Furthermore, we combined automated sample processing with shorter liquid chromatography gradients and data-independent acquisition to increase analysis throughput and enable reproducible protein quantitation across a large number of samples. We successfully applied this workflow to optimise the detection of proteasome substrates by proximity-dependent labelling.

## Introduction

Proximity dependent biotinylation is a well-established method to study protein-protein interactions in cells ^1–3^ and it is widely regarded as a complementary method to affinity purification based methods^4^. Here, a promiscuous biotin ligase is tagged to the target protein. Every protein in its close vicinity will be biotinylated and can be enriched via a subsequent pulldown with streptavidin or related affinity reagents. Since the biotinylation reaction takes place in the cell, not only stable protein-protein interactions such as protein complexes ^5^ can be detected, but also more transient interactions, e.g., enzyme-substrate interactions ^6^, impact of posttranslational modifications ^7,8^ and compartmentalization ^2,9–11^ can be monitored with this type of approach.

The strong affinity of biotin to streptavidin (K_d_=∼ 1-10*10^−15^) allows for very efficient and stringent enrichment of labelled proteins ^12^. However, in turn, the elution of biotinylated proteins from streptavidin beads can be inefficient. Instead of breaking the biotin-streptavidin interaction, most current protocols use on-bead enzymatic digestion (typically with trypsin) to recover peptides derived from the captured biotinylated proteins. This strategy allows the identification of both biotinylated proteins and their interactors. However, it suffers from two limitations. First, streptavidin can also be digested by trypsin leading to strong contamination of highly abundant streptavidin-derived peptides. Second, the fraction of biotinylated peptides recovered is typically low because they remain bound to streptavidin after digestion. The direct identification of biotinylated peptides is desirable because it enhances the confidence of the candidate protein detection and it provides structural information for direct protein-protein interactions ^13^. Therefore, different strategies have been developed to improve the detection of biotinylation sites using specific antibodies ^13,14^, modified streptavidin or (cleavable) biotin versions ^15–19^, or enrichment of specifically biotinylated peptides ^20^. Alternatively, the elution of biotinylated peptides from streptavidin can also be achieved by applying denaturing elution buffers featuring detergents ^21^, low pH ^22^, solvents ^23^ or a combination of those ^5^, after the on-bead digestion. An ideal workflow should enable the detection of interacting proteins by co-enrichment as well as the identification of biotinylated peptides to pinpoint more proximal (direct) interactions.

Here, we present an automated implementation of a workflow that combines (i) an improved on-bead digestion that minimises streptavidin contamination ^24^; (ii) a second elution step designed to enhance the recovery of biotinylated peptides ^5^ using a denaturing buffer; and (iii) label-free Data Independent Acquisition (DIA) for peptide identification and quantification (Figure 1). The new workflow enables processing of 96 samples in parallel and reduces the mass spectrometry analysis time by employing shorter chromatographic gradients. Importantly, our workflow yields more consistent protein quantification across replicates, enables more robust detection of biotinylated peptides, and provides better signal-to-noise ratio thanks to a reduced unspecific binding. Finally, the higher efficiency of the automated workflow enables input material to be reduced up to 5 fold compared to its manual version.

**Figure 1:**
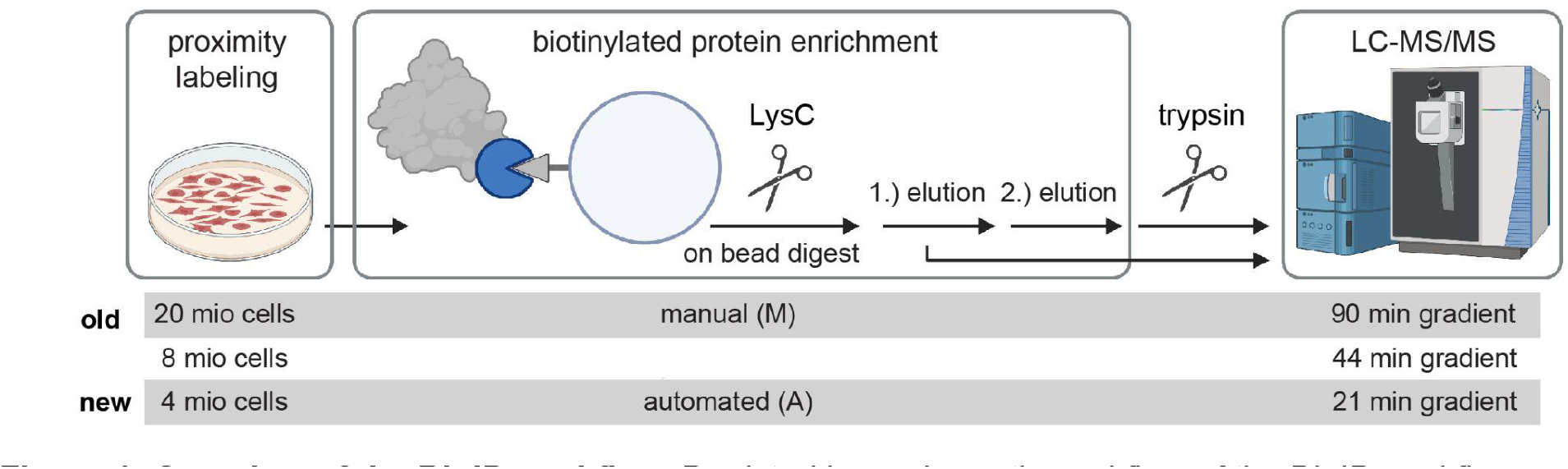
Overview of the BioID workflow. Depicted is a schematic workflow of the BioID workflow showing the different parameters tested such as cell input, enrichment procedure, and gradients of the LC-MS/MS analysis. The figure has been created with biorender.

### Experimental section

#### Generation of stable fusion protein cell lines

As described in Bartolome et al. ^24^,a cell line expressing mTurbo-PSMD3 was generated using FlpIn T-REx 293 cells (Thermo Fisher Scientific, R78007). The parental cell line was maintained in the presence of Zeocin^™^ (100 µg/ml, Thermo Fisher Scientific) and Blasticidin (15 µg/ml, Thermo Fisher Scientific). After transfection, cells expressing the constructs were selected using Blasticidin (15 µg/ml) and Hygromycin B (100 µg/ml, Thermo Fisher Scientific). PSMA4-BirA*, BirA*, PSMA4-mTurbo, or mTurbo expressing cell lines have been described in Bartolome et al. 2023. All cell lines were grown at 37°C, 5% CO_2_ and 95% humidity in Dulbecco’s modified Eagle’s medium (DMEM, Sigma Aldrich) high glucose 4.5 g/l supplemented with 10% (v/v) heat-inactivated Fetal Bovine Serum (Thermo Fisher Scientific) and 2 mM L-Glutamine (Sigma Aldrich).

#### Expression of fusion proteins

Fusion protein expression was induced with 1 µg/µl tetracycline for 4 days. For cells expressing PSMA4-BirA* or BirA*, biotin (final concentration: 50 µM, Sigma Aldrich) was added 24 h before harvesting. For cells expressing PSMA4-mTurbo, mTurbo-PSMD3 or mTurbo, biotin (50 µM) was added 2 h before harvesting. The proteasome inhibitor MG132 (Sigma Aldrich) was added to a final concentration of 20 µM 4 h before harvesting. The cells were harvested with 0.05% trypsin (Thermo Fisher Scientific) and washed with PBS three times. Cell pellets were frozen until further usage.

#### Enrichment and digest of biotinylated proteins on AssayMap Bravo

For each replicate 4, 8 or 20 Mio cells (corresponding to approximately 0.4, 0.8 or 2 mg protein) were resuspended in 250, 500 µl or 1 ml lysis buffer (50 mM Tris pH 7.5; 150 mM NaCl; 1 mM EDTA; 1mM EGTA; 1% (v/v) Triton 0; 1 mg/ml aprotinin (Carl Roth); 0.5 mg/ml leupeptin (Carl Roth); 250 U turbonuclease (MoBiTec GmbH); 0.1% (w/v) SDS), respectively, and incubated for 1 h at 4°C. For each step a modified version of the On-Cartridge protocol was used. For the acetylation of the streptavidin cartridges and the subsequent loading of the lysates, the protocol was modified as follows: Cartridges were equilibrated with 200 µl PBS (10 µl/min). For each replicate 50 µl 10 mM sulfo-NHS acetate was used. Reaction was set to 6 µl volume, 25°C for 30 min. Reaction Chase was 100 µl with a flow rate of 10 µl/min. Prior to loading cartridges were equilibrated using Internal Cartridge Wash 1, with 200 µl of lysis buffer (20 µl/min). Before and after equilibration, Cup Wash was used with default settings. Samples were loaded with a speed of 10 µl/min. Before LysC digestion cartridges were washed once with 200 µl lysis buffer and two times with 250 µl 50 mM ammonium bicarbonate (AmBic) (10 µl/min). For each digest 0.5 µg LysC (Cell Signaling) was added to 30 µl 50 mM AmBic. Reaction was set to 6 µl volume, 45°C, 60 min with no Reaction Chase. Peptides were eluted in two steps (1x 25 µl and 1x 50 µl) 50 mM AmBic (no internal cup wash, 10 µl/min). Trypsin (0.5 µg, Promega) was added to elution and incubated at 37°C overnight. Biotinylated peptides were eluted with two times 15 µl 10% TFA in acetonitrile (90 µl/min). Syringe was washed with 150 µl 20% acetonitrile. Biotinylated peptide elution was dried and reconstituted in 50 µl 50 mM HEPES. The pH was tested and adjusted with sodium hydroxide to pH 6 −8.0. Eluate was digested with 0.5 µg trypsin at 37°C overnight. Both elutions were cleaned up using Waters Oasis® HLB μElution Plate 30 μm (Waters) according to the manufacturer’s instructions.

#### Manual enrichment and digestion of biotinylated proteins

The protocol was used as described in Bartolome et al ^24^. In short, 20 Mio cells were resuspended in 4.75 ml lysis buffer (see above) and incubated for 1 h at 4°C. Streptavidin Sepharose High Performance (GE Healthcare) was acetylated by the addition of 10 mM sulfo-NHS acetate (Thermo Fisher Scientific) for 30 min twice. For each lysate 80 µl of equilibrated beads were used. Beads were washed 5 times with 600 µl 50 mM AmBic. On-bead digest was performed with 200 µl LysC (5 ng/µl) at 37°C overnight. First elution step was achieved using two times 150 µl of 50 mM AmBic. After pooling both fractions, peptides were further digested by adding 1 µg trypsin and incubating at 37°C for 3 h. Biotinylated peptides were eluted using two times 150 µl 20% TFA (Biosolve) in acetonitrile (Biosolve). Both fractions were pooled and neutralised to pH 8.0 by adding 50 µl 200 mM HEPES and sodium hydroxide as necessary. Peptides were digested through the addition of 1 µg trypsin at 37°C for 3 h. Both elutions were desalted using Waters Oasis® HLB μElution Plate 30 μm (Waters) according to the manufacturer’s instructions.

#### Immunoblot

Lysates (10 μg protein) was separated on 4%–20% Mini-PROTEAN® TGX Precast Protein Gels (Biorad) and blotted onto a Roti®-NC transfer membrane (Carl Roth). Proteins were visualized with Ponceau S staining before incubation with 3% BSA (Thermo Fisher) in TBST for 1 h at room temperature. Membranes were then incubated with the Streptavidin-HRP (1:20000, Abcam ab7403) for 1 h at room temperature. After washing with TBST the membranes were incubated with Pierce ECL Western Blotting Substrate (Thermo Fisher Scientific) and detection was carried out using a ChemiDocTM XRS+ Imaging system (Biorad).

#### LC-MS analysis

For in-depth proteomics analysis, approximately 1μg of reconstituted peptides were separated using a nanoAcquity UPLC (Waters) coupled online to the MS. Peptide mixtures were separated in trap/elute mode, using a trapping (Waters nanoEase M/Z Symmetry C18, 5μm, 180 μm x 20 mm) and an analytical column (Waters nanoEase M/Z Peptide C18, 1.7μm, 75μm x 250mm). The outlet of the analytical column was coupled directly to an Orbitrap Fusion Lumos mass spectrometer (Thermo Fisher Scientific) using the Proxeon nanospray source. Solvent A was water, 0.1% formic acid and solvent B was acetonitrile, 0.1% formic acid. The samples were loaded with a constant flow of solvent A, at 5 μL/min onto the trapping column. Trapping time was 6 min. Peptides were eluted via the analytical column with a constant flow of 300 nL/min. During the elution step, the percentage of solvent B increased in a nonlinear fashion from 0% to 40% in 90 min. Total runtime was 115 min, including cleanup and column re-equilibration. The peptides were introduced into the mass spectrometer via a Pico-Tip Emitter 360 µm OD x 20 µm ID; 10 µm tip (New Objective) and a spray voltage of 2.2 kV was applied. The capillary temperature was set at 300 °C. The RF lens was set to 30%. Full scan MS spectra with mass range 350-1650 m/z were acquired in profile mode in the Orbitrap with resolution of 120,000 FWHM. The filling time was set at maximum of 20 ms with an AGC target of 5 x 10^5^ ions. DIA scans were acquired with 34 mass window segments of differing widths across the MS1 mass range. The HCD collision energy was set to 30%. MS/MS scan resolution in the Orbitrap was set to 30,000 FWHM with a fixed first mass of 200m/z after accumulation of 1x 10^6^ ions or after filling time of 70ms (whichever occurred first). Data were acquired in profile mode. For data acquisition and processing Tune version 3.5 and Xcalibur 4.5 were employed.

For high-throughput analysis on Evosep, the samples were loaded on Evotips according to the manufacturer’s instructions. In short, Evotips were first washed with Evosep buffer B (0.1% formic acid in acetonitrile), conditioned with 100% isopropanol and equilibrated with Evosep buffer A (0.1% acetonitrile). Afterwards samples were loaded on the Evotips and washed with Evosep buffer A. The loaded Evotips were topped up with buffer A and stored until the measurement. Peptides were separated using the Evosep One system (Evosep, Odense, Denmark) equipped either with a 8 cm x 150 μm i.d. packed with 1.5 μm Reprosil-Pur C18 beads column (Evosep Performance, EV-1109, PepSep) for the pre-programmed proprietary Evosep gradient of 21 min (60 samples per day, 60SPD), or with a 15 cm x 150 μm i.d. packed with 1.9 μm Reprosil-Pur C18 beads column (Evosep Endurance, EV-1106, PepSep) for the re-programmed proprietary Evosep gradient of 44 min (30 samples per day, 30SPD). Solvent A was water and 0.1% formic acid and solvent B was acetonitrile and 0.1% formic acid. The LC was coupled to an Orbitrap Exploris 480 (Thermo Fisher Scientific) using the Proxeon nanospray source. The peptides were introduced into the mass spectrometer via a PepSep Emitter 360-μm outer diameter × 20-μm inner diameter, heated at 300 °C, and a spray voltage of 2 kV was applied. The radio frequency ion funnel was set to 30%. For DIA data acquisition of the 60SPD method, full scan mass spectrometry (MS) spectra with mass range 350–1650 m/z were acquired in profile mode in the Orbitrap with resolution of 120,000 FWHM. The default charge state was set to 2+. The filling time was set at a maximum of 45 ms with a limitation of 3 × 106 ions. DIA scans were acquired with 35 mass window segments of differing widths across the MS1 mass range. Higher collisional dissociation fragmentation of 30%) was applied and MS/MS spectra were acquired with a resolution of 15,000 FWHM with a fixed first mass of 200 m/z after accumulation of 1 × 106 ions or after filling time of 37 ms (whichever occurred first). Data were acquired in profile mode. For DIA data acquisition of the 30SPD method, full scan mass spectrometry (MS) spectra were acquired as above, with the only difference of the filling time set at at a maximum of 60 ms with a limitation of 3 × 106 ions. DIA scans were acquired with 40 mass window segments of differing widths across the MS1 mass range. Higher collisional dissociation fragmentation of 30% was applied and MS/MS spectra were acquired with a resolution of 30,000 FWHM with a fixed first mass of 200 m/z after accumulation of 1 × 106 ions or after filling time of 45 ms (whichever occurred first). Data were acquired in profile mode. For data acquisition and processing of the raw data Xcalibur 4.5 (Thermo) and Tune version 4.0 were used.

#### Data Analysis

DIA raw data were analyzed using the directDIA pipeline in Spectronaut (PSMA4-miniTurbo samples with v.18, everything else v.17, Biognosys). The data were searched against a specific species (Homo sapiens, 20,816 entries) and a contaminants (247 entries) Swissprot database. The data were searched with the following modifications: Oxidation (M), Acetyl (Protein N-term) and Biotin_K. A maximum of 2 missed cleavages for trypsin and 5 variable modifications were allowed. The identifications were filtered to satisfy FDR of 1% on peptide and protein level. Relative quantification was performed in Spectronaut using LFQ QUANT 2.0 method with Global Normalization, precursor filtering percentile using fraction 0.2 and global imputation. The data (candidate table) and data reports (protein quantities) were then exported and further data analyses and visualization were performed with Rstudio using in-house pipelines and scripts. The significance of the increase of the fold change from proteasome & associates was calculated using the Wilcoxon Rank Sum Test.

To identify ProteasomeID enriched proteins, we trained a logistic regression binary classifier. The classifier was trained using known proteasome interactors as positive class, and mitochondrial matrix proteins as negative. Mitochondrial matrix proteins are naturally biotinylated, but they are not expected to interact directly with the proteasome under homeostatic conditions. To distinguish between these two classes, we performed prediction using an enrichment score derived from multiplying the average log2 ratio and the negative logarithm of the q-value obtained from a differential protein abundance analysis performed against the BirA* control line. Before analysis, any missing data points were removed from the dataset. To assess the performance of the binary classifier, and optimize its parameters, a 10-fold cross-validation approach was adopted. The dataset was randomly partitioned into ten subsets, with nine subsets used for training and one subset for validation in each iteration. This process was repeated thirty times, and results were averaged using the mean value to ensure the robustness of the results. Logistic regression was employed as the classification method using the caret package in R ^25^. To determine an optimal threshold for classification, the false positive rate (FPR) was set at 0.05. The threshold yielding an FPR closest to the target value of 0.05 was selected as the final classification threshold. Following model training and threshold selection, the classifier was applied to predict the class labels of additional proteins not used in the training process. The enrichment score and class labels for the new data were provided as input to the trained model. All statistical analyses were performed using R version 4.1.3. The pROC ^26^ and caret ^25^ packages were employed for ROC analysis and logistic regression, respectively. The F1 score used to compare conditions was calculated as the harmonic mean of precision and recall: TP/(2TP+FP+FN), TP: true positive, FP: false positive, FN: false negative.

#### Data availability

Mass spectrometry proteomics data have been deposited to ProteomeXchange Consortium via the MassIVE partner repository and they are accessible with the identifier MSV000092703.

## Results

### Implementation of an automated workflow for high-throughput BioID

In this study, we used HEK293T cells expressing the PSMA4/alpha3 subunit of the human proteasome tagged with the promiscuous biotin ligase BirA*, as a proxy for method development. We have previously characterised this cell line both in terms of enrichment efficiency and known associated proteins pulldown ^24^. First, we adapted a sample preparation workflow for enrichment and digestion of biotinylated proteins ^24^, and implemented it on the liquid handler Bravo AssayMAP (Figure 2a and Supplemental Figure 1a). The workflow has been optimised for parameters such as lysate concentration, beads pretreatment, and digest time. Mass spectrometry analysis of peptides obtained by the automated workflow showed clear separation of samples from different experimental groups by principal component analysis (PCA) (Figure 2a), and the expected enrichment of proteasome subunits and associated proteins in the comparison of PSMA4-BirA* vs. BirA* control (Supplemental Figure 1b). We also confirmed, as previously shown in many applications, that acetylation of the streptavidin beads prior to loading reduces streptavidin contamination without impacting biotin binding to streptavidin (Supplemental Figure 1c). We also tested several washing and priming conditions as suggested by the manufacturer. We decided to omit the priming step with 1% FA as it caused an increase of singly charged ions in the samples. Moreover, washing the columns with a similar bead volume as used with the manual protocol was not sufficient to remove detergents (Supplemental Figure 1d) and impacted the enrichment (Supplementary figure 1b). After removal of the detergents through increased washing, the proteasome and its associated proteins clearly separated from the other proteins showing a higher log2 ratio as well a high protein group quantity. Of note, most of the lid components of the 19S regulatory particle of the proteasome were not enriched (Supplemental Figure 1b), likely due to an artefact of the sample processing. The lysates were frozen before enrichment and this might have led to a partial disassembly of the proteasome ^27^. Most of the lid proteins are in fact not close enough to PSMA4 to achieve direct biotinylation ^24^, and were therefore not enriched directly during the streptavidin pulldown. In the subsequent analyses, the lid proteins were omitted from the list of proteasome and associated proteins to account for this artefact and do not bias the comparison of different methods.

**Figure 2:**
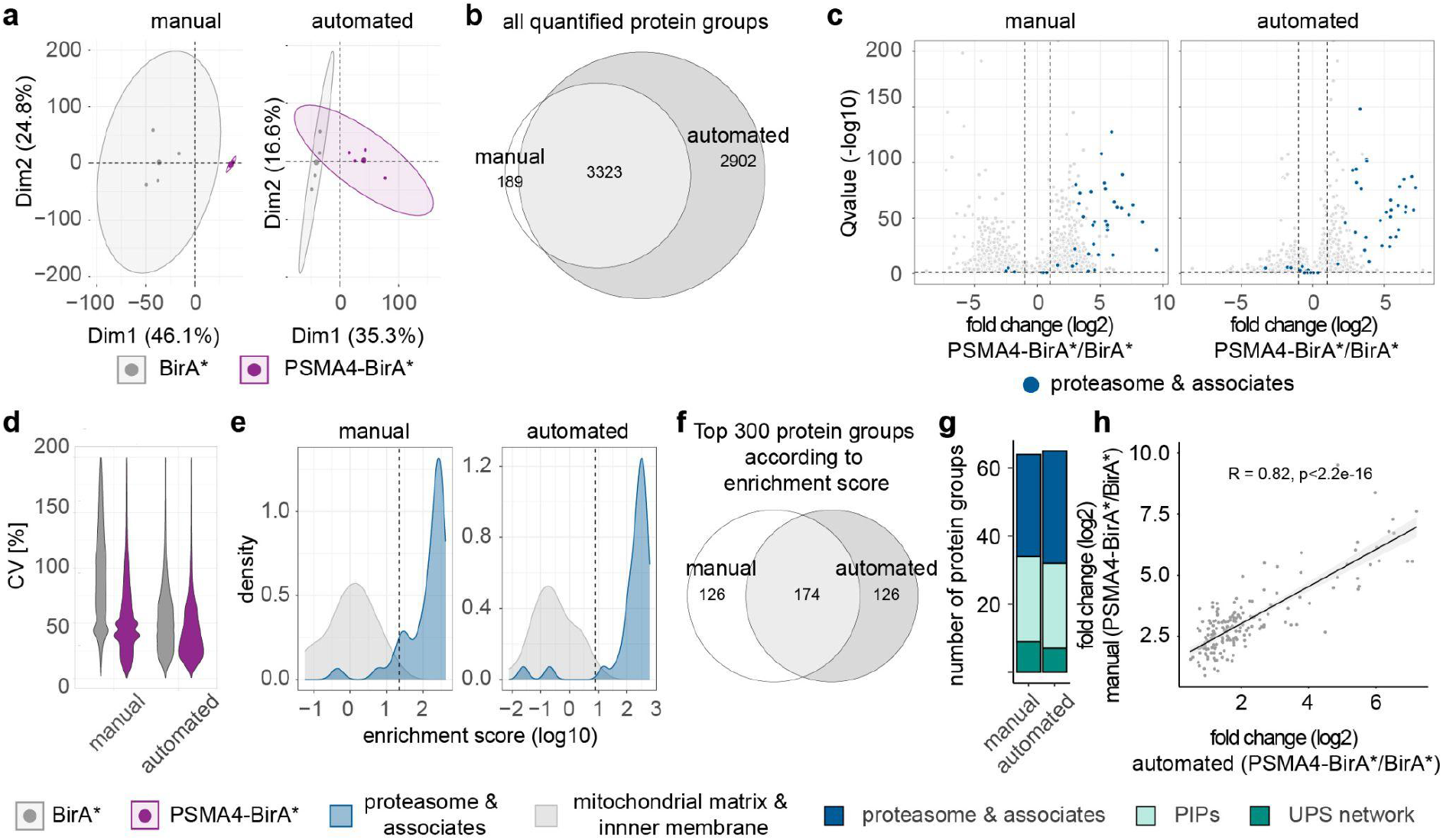
Comparison of manual and automated BioID workflows. A streptavidin pulldown of HEK293 expressing either PSMA-BirA* or BirA* was performed using either the manual or automated workflow. Here, we analysed the first elution of the pulldowns. Data shown are from four experiments. (a) Principal component analysis of biotin-enriched samples from either the manual or automated workflow. Smaller dots represent replicates, while larger dots are the centroids of each sample group. Ellipses represent 95% confidence intervals. (b) Venn diagram for all quantified protein groups using either the manual (white) or the automated (dark grey) workflow. (c) Volcano plots for quantified protein groups for both workflows. Proteasome subunits and known associated proteins are highlighted in dark blue. Dashed lines were set at log2 fold change > 1 and Q value < 0.05. n = 4. (d) Distribution of the coefficient of variation (CV) of both enrichment strategies for BirA* (grey) and PSMA4-BirA* (purple) expressing cells. (e) Distribution of true positive (proteasome subunits and associates, blue) and true negative (mitochondrial matrix and inner membrane proteins, grey) proteins according to their enrichment score. Dashed line marks the calculated cutoff for a true positive rate (FPR) < 0.05. (f) Venn Diagram for the top 300 protein groups according to the enrichment score comparing the manual (white) with the automated (grey) workflow. (g) Number of proteins in the top 300 that could be identified as either proteasome subunits and associated proteins (dark blue), known proteasome interacting proteins (PIPs, light turquoise)) or the ubiquitin-proteasome system (UPS, dark turquoise) network. (h) Correlation of the average log2 ratio of the proteins shared in the top 300 proteins.

### Comparison of manual and automated BioID workflows

After optimization of the protocol, we compared our automated workflow to the manual version. Almost 95% of the quantified proteins from the manual protocol were shared with the automated workflow (Figure 2b). Most notably, more protein groups were quantified using the automated workflow including 2902 protein groups that were not identified in the manual version. For both protocols, the proteasome and associated proteins were significantly enriched compared to BirA* control. Interestingly, the median average log2 ratio of proteins showing higher enrichment in BirA* control was higher using the manual (2.25) than the automated workflow (0.72) (Figure 2c). This might indicate a higher level of unspecific binding of proteins to the matrix as the bead volume is larger in the manual protocol (80 µl beads) than for the automated workflow (5 µl beads). This observation is supported by the fact that the BirA* samples from the manual workflow showed the highest coefficient of variation (CVs) (Figure 2d). As expected, the CVs from the PSMA4-BirA* samples were lower for both workflows. To better assess the signal-to-noise ratio, we analysed the separation between naturally biotinylated mitochondrial proteins (true negatives) and proteasome subunits and assembly factors (true positives) using a logistic regression classifier that we previously developed ^24^. Briefly, we calculated an “enrichment score”, which combines the negative logarithm of the Q-value with the average log2 ratio, and ranked the proteins according to their enrichment score. We found a clear separation between the true negatives and true positives for both the manual and automated workflow (Figure 2e). In order to compare the performance of the protocols, we used the F1 score, which combines the accuracy of positively identified proteins with the number of retrieved true positives (see Methods). We found that the overlap between the distributions of enrichment scores for the positive and negative sets was larger for the manual (F1 score: 0.903) than the automated (F1 score: 0.955) protocol, likely due to the higher unspecific binding. Taken together, these observations indicate that the reduced volume of beads in the automated workflow decreased the background binding and consequently improved the signal-to-noise ratio.

We then specifically analysed the top 300 proteins according to the enrichment score as these should contain most of the known proteasome subunits and interactors. More than half of the proteins (174 protein groups) were found in both workflows (Figure 2f). For both workflows similar numbers of proteasome subunits & associates (27 manual, 33 automated), proteasome-interacting proteins PIPs (9 manual, 7 automated) and proteins from the ubiquitin-proteasome system UPS (25 manual, 30 automated) were found in the top 300 proteins (Figure 2g). Moreover, the average log2 ratio from the top 300 proteins between the manual and the automated workflow was highly correlated (Figure 2h).

In summary, our data indicate that the new workflow on the Bravo AssayMAP performs as well as the manual workflow in terms of enrichment of known proteasome subunits and interactors in the top 300 proteins, while reducing background binding and increasing the quantification of protein groups overall. Importantly, the automated workflow enables parallel processing of up to 96 BioID samples in just 2 days, while the normal protocol allows to process maximum up to 24 samples at once in the same time

### Influence of starting material and gradient length

We reasoned that the higher efficiency and improved signal-to-noise of the automated workflow could enable BioID analysis from lower input material and with faster analysis. Therefore, we used our automated workflow to evaluate the impact of sample input (number of cells used for each enrichment) and gradient length of the LC-MS/MS analysis. We compared 8 combinations of input material and gradient length for a total of 128 MS runs. We found no clear correlation between the amounts of input tested and the identification of precursors, peptides or protein groups (Figure 3a, Suppl. Figure 2a-c). However, higher cell input correlated with an enhanced enrichment of proteasomal proteins, which was shown through a significantly higher average log2-fold change for proteasomal proteins between 20 Mio and 4 Mio cell input (Figure 3b). Consistently, the separation of true negatives and true positives using our enrichment score was better with increasing cell input, as reflected by F1 scores (4 Mio: 0.861, 8 Mio: 0.870, 20 Mio: 0.955) (Figure 3c). Despite this, when considering the top 300 enriched proteins, we found no major difference in the recovery of proteasome subunits and interactors, previously reported proteasome-interacting proteins (PIPs) ^28–30^ and members of the UPS network proteins (Figure 3d).

**Figure 3:**
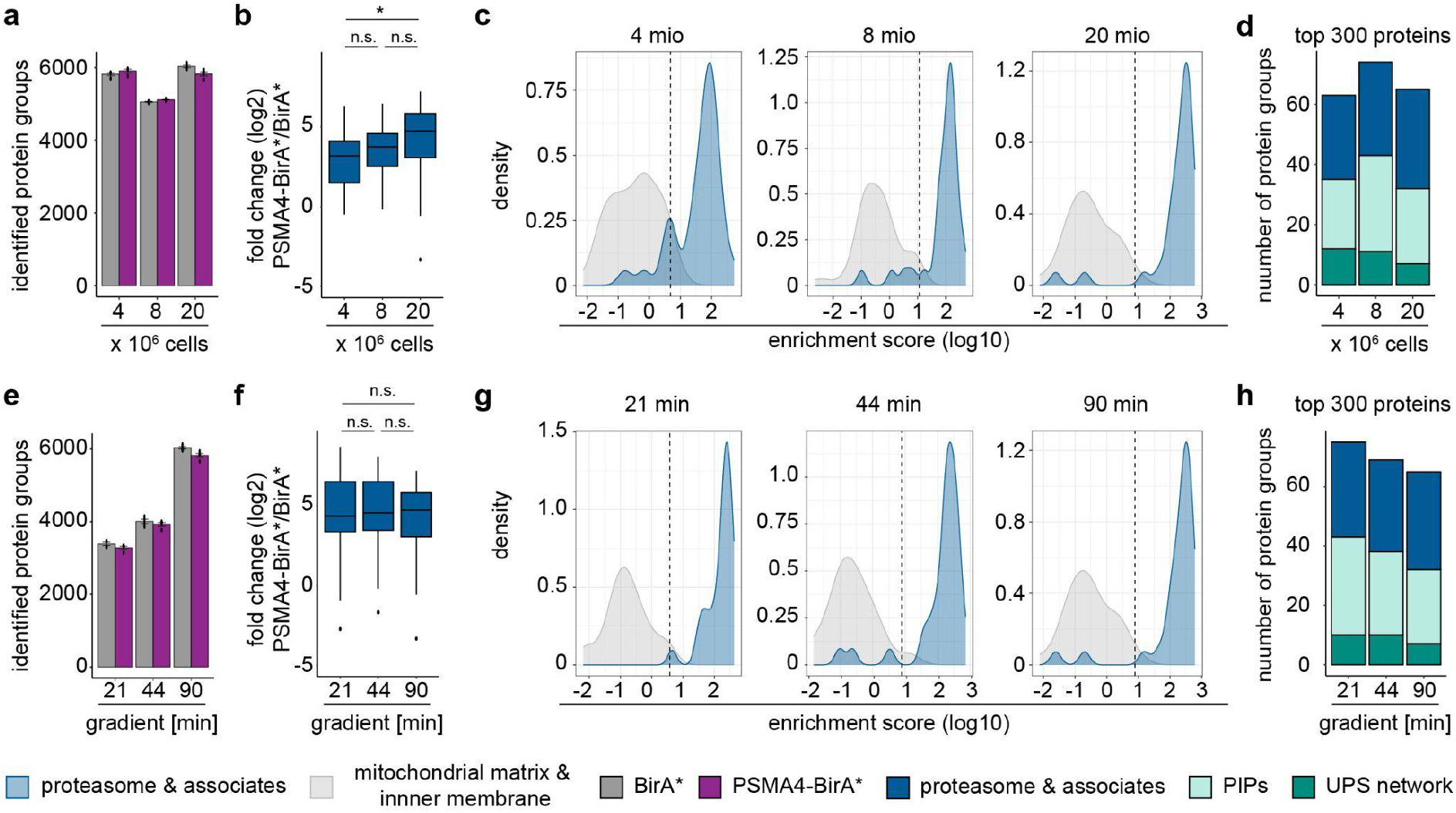
Influence of starting material and gradient length. To test the influence of input and gradient length of LC-MS/MS analysis we first compared three different cell numbers (4, 8 and 20 Mio) for the automated streptavidin enrichment with the same gradient (90 min gradient). Afterwards, we kept the input amount stable, while varying the gradient length (21 min, 44 min and 90 min). Only the first elution has been analysed in this figure. All data shown are from 4 independent experiments. (a) Number of identified protein groups of proteins separated by cell lines, BirA* (grey) and PSMA4-BirA* (purple) expressing, and input amount. (b) Higher enrichment of proteasome subunits and associates in PSMA4-BirA* samples depending on cell input. *p<0.05, Wilcoxon Rank Sum Test. (c) Distribution of true positive (blue) and true negative (grey) proteins according to the enrichment score. Dashed line marks the calculated cutoff for a true positive rate (FPR) < 0.05. (d) Number of proteins in the top 300 that could be identified as either a proteasome subunits and associated proteins (dark blue), known PIPs (light turquoise) or member of the UPS network (dark turquoise) network. (e) Number of identified protein groups of proteins separated by cell lines, BirA* and PSMA4-BirA* expressing, and gradient length. (f) Enrichment of proteasome subunits and associates in PSMA4-BirA* samples independent of gradient length. *p<0.05, Wilcoxon Rank Sum Test. (g) Distribution of true positive and true negative proteins according to the enrichment score. Dashed line marks the calculated cutoff for a true positive rate (FPR) < 0.05. (h) Number of proteins in the top 300 that could be identified as either a proteasome subunit and associated proteins, known PIPs or the UPS network.

Next, we evaluated the impact of the LC gradient length keeping cell input fixed at 20 Mio. As expected, gradient length affected the overall identification of precursors, peptides or protein groups, with more identifications in the longest gradient (Figure 3e, Suppl. Figure 2a-c). However, it did not influence the relative enrichment of proteasomal and related proteins, as shown by comparison of average log2 fold changes (Figure 3f), and no major difference in the separation between enrichment scores of true negatives and true positives (F1 scores for 21 min: 0.923, 44 min: 0.938, 90 min: 0.955) (Figure 3g). Interestingly, we observed a higher recovery of proteasome-related proteins among the top 300 enriched proteins using shorter gradients (Figure 3h). From these results, we conclude that shorter gradients down to 21 min can be used in combination with our automated enrichment workflow, enabling higher throughput without negative impact on the quality of the analysis.

### Automated workflow to increase identification of biotinylation sites

Finally, we evaluated the enrichment of biotinylated peptides using the second elutions from 20 Mio cells input analysed using a 90 min gradient. Using direct DIA analysis, we could show that the automated workflow enabled the detection of almost twice the number of biotinylated peptides in comparison to its manual counterpart (678 against 351) (Figure 4a). As expected, the log2 fold changes obtained for the biotinylated peptides correlated with the ones obtained for the corresponding proteins from the on-bead digestion fraction, with proteasome subunits and related proteins displaying the strongest enrichment (Figure 4b). More biotinylated proteins from the automated (80 protein groups) compared to the manual (44 protein groups) workflow were assigned to proteasome and associates, PIPs or the UPS network (Figure 4c). These results show that the automated workflow improves detection of biotinylated peptides and therefore the identification of direct interactors.

**Figure 4:**
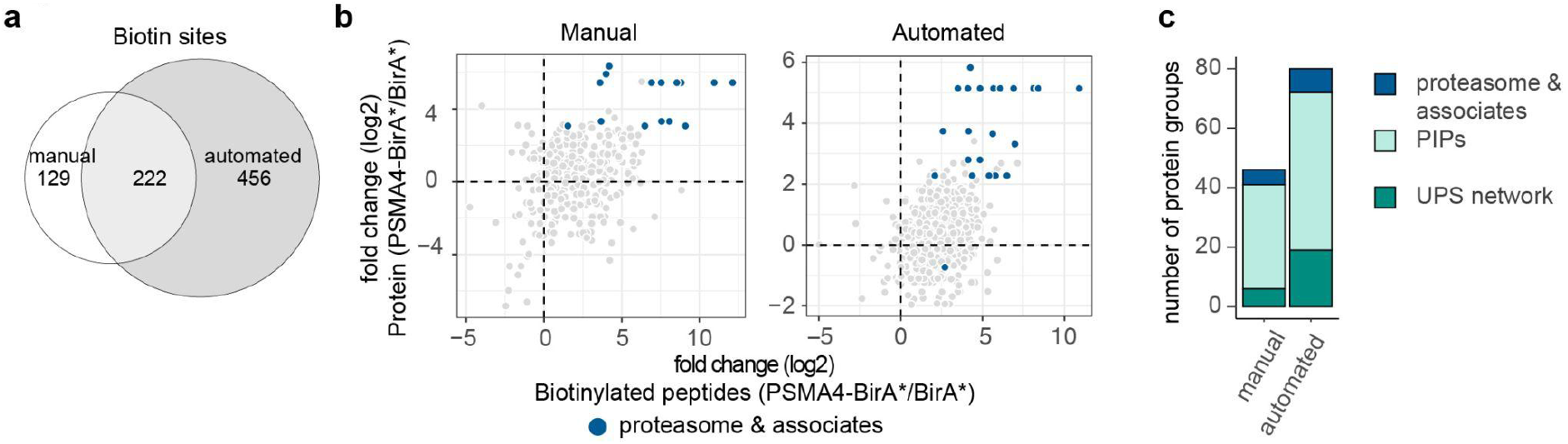
Automated workflow increases the identification of biotinylation sites. A streptavidin pulldown of HEK293 expressing either PSMA-BirA* or BirA* was performed using either the manual or automated workflow. Here, we analysed the second elution of the pulldowns containing most of the biotinylated peptides. Data shown are from four experiments. (a) Venn diagram comparing identified biotinylated peptides using the manual (white) with the automated (dark grey) workflow. (b) Plotting the average log2-fold enrichment of proteins against the average log2-fold enrichment of biotinylated peptides. Proteasome subunits and known associated proteins are highlighted in dark blue. (c) Number of identified biotinylated proteins of the proteasome and associates, PIPs or UPS network in the automated vs manual workflow.

### Application of automated BioID to improve the detection of proteasome substrates

Next, we wanted to demonstrate the application of the optimised workflow for improving the detection of proteasome substrates by proximity-dependent labelling, an approach that we have previously developed using the manual BioID workflow ^24^. Therefore, we tagged two subunits of the proteasome with miniTurbo, as it has been shown to biotinylate more rapidly ^31^. Furthermore, the proteasome was inhibited with MG132 to prolong the duration of the interaction between the proteasome and its substrates and reduce non-tryptic peptide generation, as previously shown ^24^. To test whether a subunit from the 19S would improve detection of substrates, PSMD3 was chosen in addition to the previously characterised 20S subunit PSMA4 (Figure 5a, Supplemental Figure 3a-b). Proteins that were found enriched from lysates of cells expressing PSMA4-miniTurbo or miniTurbo-PSMD3 treated with MG132, but not in samples treated with DMSO were defined as potential substrates (Figure 5b-c). While we detected 206 potential substrates with PSMA4-miniTurbo, almost double the amount of potential substrates (437) was identified with miniTurbo-PSMD3 (Figure 5d). Out of these more than half have been previously reported to display an increased ubiquitylation in response to proteasome inhibition ^32^. Most of the known substrates found with PSMA4-miniTurbo were also found with miniTurbo-PSMD3 (Figure 5e). However, the majority of these known substrates from our miniTurbo-PSMD3 analysis were found as significantly enriched exclusively with this construct. Taken together, these results suggest that miniTurbo-PSMD3 construct is better for detection of proteasome substrates by proximity dependent labelling than PSMA4-miniTurbo.

**Figure 5:**
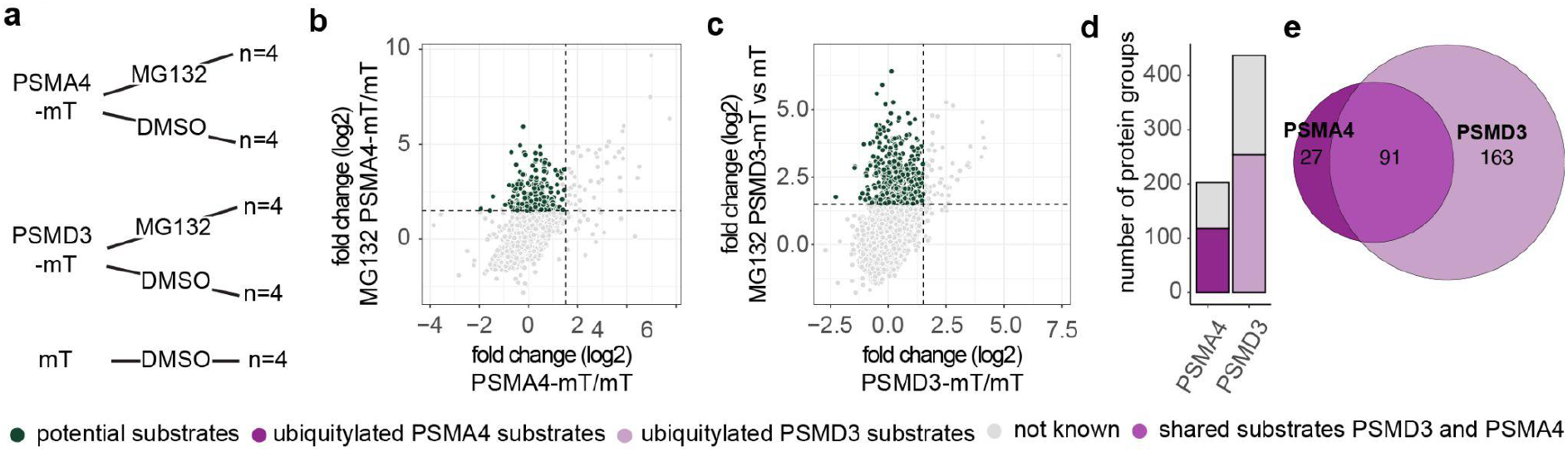
Tagging different proteasome subunits to identify proteasome substrates. (a) Workflow to identify the best construct for substrate detection. Two proteasome subunits from either the 19S (PSMD3) or 20S (PSMA4) complex were fused to miniTurbo. To enhance detection of substrates, proteasome inhibitor MG132 was added. Data shown here are from 4 independent experiments. Differential expression of (b) PSMA4-miniTurbo or (c) miniTurbo-PSMD3 compared to miniTurbo with MG132 or vehicle control was plotted against each other. Proteins enriched after inhibition (AVG Log2 > 1.5, Q-value < 0.05), but not with the vehicle control (AVG Log2 <1.5, Q-value < 0.05) were defined as potential substrates (dark green). (d) Number of potential substrates that have been shown by Trullson et al to increase ubiquitylation after MG132 treatment. (e) Venn diagram comparing ubiquitylated protein groups identified using PSMA4-miniTurbo or miniTurbo-PSMD3.

## Discussion

During recent years, proximity dependent biotinylation has become the method of election to study protein-protein interactions and it has been vastly developed and optimised for many different applications including studies of protein complexes and cell compartmentalisation ^5,9,33^. The biotinylation reaction has been improved by using different different promoters or antibody-based delivery of the biotin ligase ^34–36^. Some recent developments include the combination of mass spectrometry for posttranslational modifications (PTMs) with proximity labelling methods to elucidate the role of PTMs on the localisation of proteins ^7,8^.

The increasing complexity of experimental designs for this type of approaches, which often require multiple controls to enable correct interpretation of the results, motivated us to develop a robust and higher throughput workflow for sample preparation and MS analysis. By combining automated sample preparation on a robot with short gradient LC-MS methods, our pipeline reduces the sample processing and measuring time up to four times compared to the manual protocol, and it enables analysis from reduced input material down to one fifth of the usual pipeline.

To our knowledge there has been only one other pre-print from the Hüttenhain lab ^37^ that describes an automated workflow for proximity dependent labelling on an alternative platform. While their manuscript optimised the parameters for the DIA acquisition, we focused more on the LC and were able to shorten the gradient to 21 min compared to their 66 min gradient. Interestingly, in both manuscripts the optimal amount was identified at around 1 mg protein or 8 Mio cells (approximately 0.8 mg). However, as we used a previously characterised construct we were able to assess that we still obtained satisfactory results from 4 Mio cells. We have not tested lower numbers of cells, thus it might be possible to obtain reasonable results from even lower input. This could be particularly relevant for applications where the input material might be limited. For instance, proximity dependent labelling has been successfully applied in organisms to study protein-protein interactions *in vivo* ^*3,38,39*^ and recently has been used to study cell-type specific proteomes in the brain ^40^. Protein complexes like the proteasome can have different compositions dependent on the cell type ^41^, and studying these complexes could give valuable insight into their cell-type specific functions. The lower sample input required by our automated BioID workflow could enable this type of studies also in less abundant cell types. However, this would need to be tested for each specific application since other factors, such as the expression level of the bait protein, might influence the yield of the streptavidin enrichment.

Finally, we also improved the detection of biotinylated peptides. The detection of direct biotinylation typically indicates that the protein was in close proximity to the bait and therefore, it likely represents a direct interaction partner. This is important as during the streptavidin pulldown not only direct interactors, but also their binding partners can be enriched. Several attempts have been made to enhance the detection of these biotinylated proteins by modifying biotin affinity reagents ^16,19^, performing pulldown on peptides (DiDBiT) ^20^ or a combination of both ^13^. We adapted the protocol developed by Bartolome et al., which has the advantage that both direct and indirect interactors are captured in the first elution step from the on-bead digestion, while biotinylated peptides are retrieved in a second elution step that is analysed separately. Because of the harsher buffer used, the second elution step is especially sensitive to variations in sample handling, e.g., contact time between elution buffer and streptavidin beads. For example, excessively long elution time can lead to denaturation of streptavidin and release of its monomers, which could negatively influence the downstream LC-MS analysis. By implementing the workflow on a liquid handler, we have reduced the variability of this step and enabled a more reproducible and deeper quantification of biotinylated peptides.

The characteristics highlighted above make our workflow suitable for most proximity labelling experiments. Furthermore, our method could be easily adapted to other protocols that rely on the enrichment of biotinylated peptides, e.g., surface proteomics or protein synthesis analysis by incorporation of amino acid analogues that can be biotinylated via click chemistry ^42–44^.

## Acknowledgments

The FLI is a member of the Leibniz Association and is financially supported by the Federal Government of Germany and the State of Thuringia. This work was also supported by the FLI Core Facilities Technology Transfer.

## Author contributions

Conceptualization: E.C., A.O, T.D, Experimental procedure: N.P., E.C., N.R., H.K., I.H., Data analysis: E.C., T.D., H.K., D.D.F., A.O., Supervision: A.O., T.D., Visualisation: E.C., H.K., A.O., T.D., writing: E.C., T.D., A.O. with input from H.K.. All authors have read and given approval to the final version of the manuscript.

## Figures

**Supplemental Figure 1:**
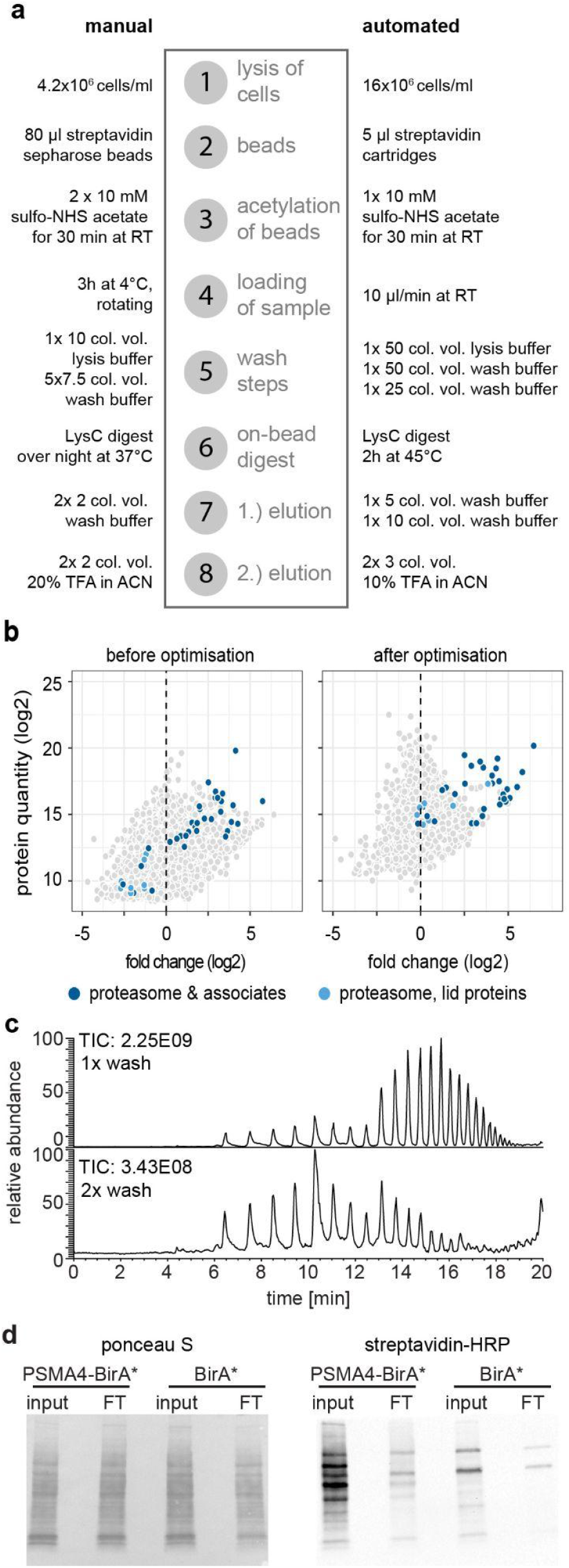
In this figure we depict the different initial washing steps that were optimised. (a) A schematic overview comparing the manual and automated workflow after optimization (b) MA plot of enriched proteins with proteasomal proteins highlighted in dark blue and proteasome lid proteins in light blue before and after washing steps were tested. (c) Chromatogram of first elution using the original (top panel) versus the optimised washing protocol (lower panel) (d) Input and flowthrough (FT) were analysed by SDS-PAGE and western blotting using streptavidin-HRP (right panel). Ponceau S was used as a loading control (left panel). HRP: horseradish peroxidase

**Supplemental Figure 2:**
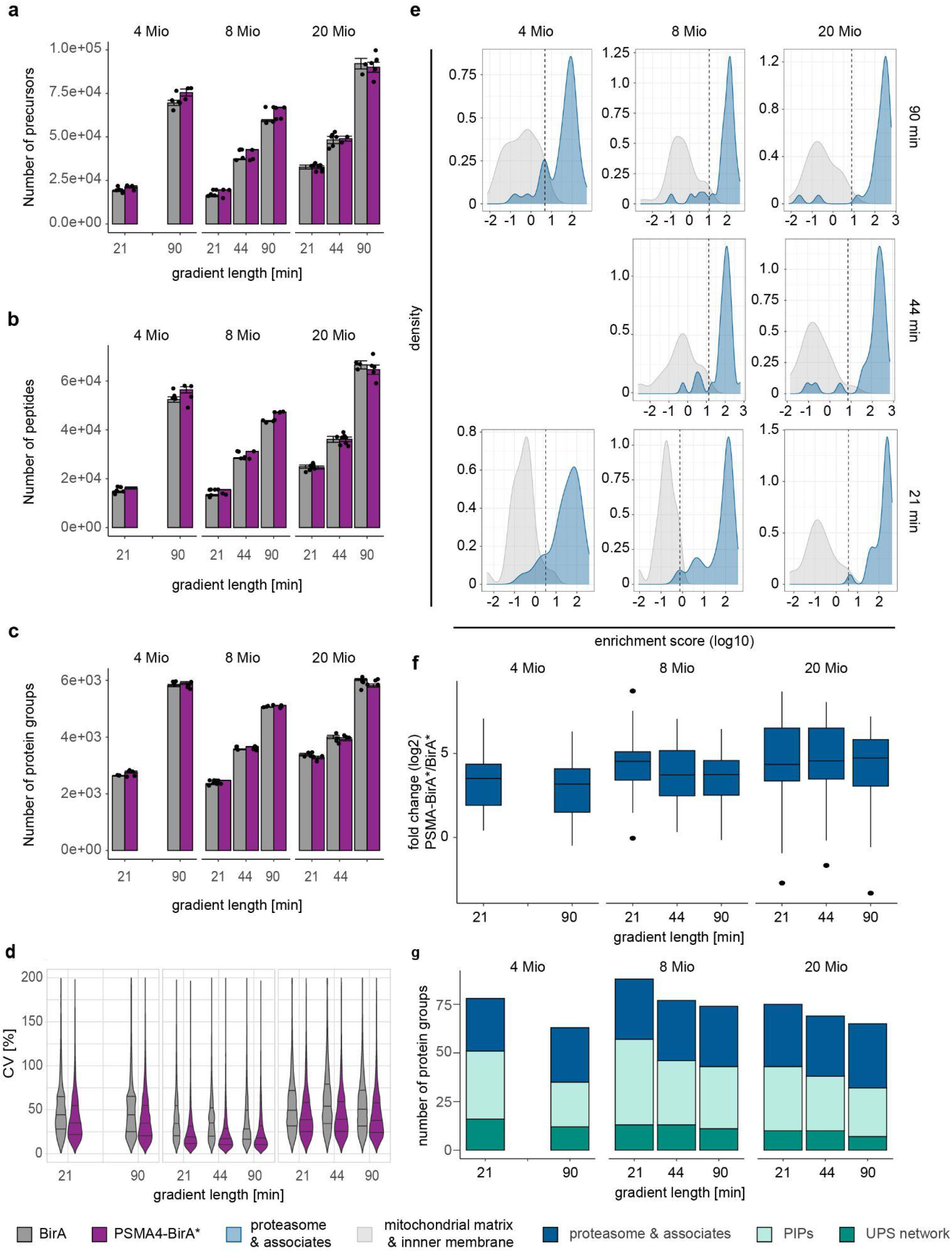
Influence of input amount and gradient length of the LC-MS/MS analysis on interactors identification. All experiments depicted in this figure show results depending on input amount (4, 8 and 20 Mio cells) and gradient length (21, 44 and 90 min). Data shown are derived from 4 independent experiments. The identification of (a) precursors, (b) peptides and (c) protein groups was impacted by the gradient, but not the cell input. For 4 and 8 Mio cells, less precursors, peptides and proteins were identified in BirA* compared to PSMA-BirA* samples. (d) Violin plot showing the distribution of CVs in BirA* and PSMA4-BirA* cell lines. (e) Distribution of true positive (blue) and true negative (grey) proteins according to the enrichment score. Dashed line marks the cutoff for a true positive rate (FPR) < 0.05. (f) Enrichment of proteasome subunits and associates in PSMA4-BirA* samples. (g) Number of proteins in the top 300 that could be identified as either a proteasome subunit and associated proteins, known PIPs or the UPS network.

**Supplemental Figure 3:**
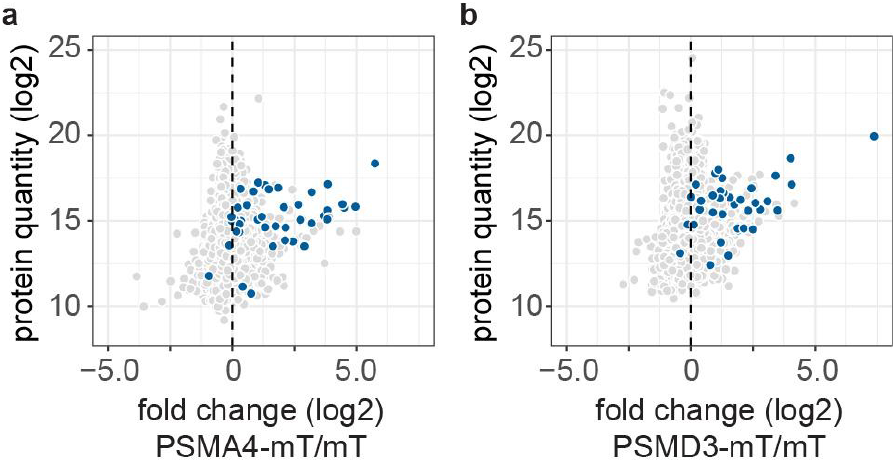
Tagging different proteasome subunits to identify proteasome subunits. Two proteasome subunits from either the 19S (PSMD3) or 20S (PSMA4) complex were fused to miniTurbo. Data shown here are from 4 independent experiments. MA plot of enriched proteins with proteasomal proteins highlighted in dark blue from either (a) PSMA4-miniTurbo or (b) PSMD3-mini-Turbo expressing cell lines.

